# Rapid Bacterial Identification through Biophysical Surface Topography Features

**DOI:** 10.64898/2026.03.12.711416

**Authors:** Adam Krueger, Bikash Bogati, Raymond Copeland, Maryam Sadat Hejri Bidgoli, Jin Zhu, David S. Weiss, Peter J. Yunker

## Abstract

Scientists have long classified organisms based on shared morphological characteristics. However, modern taxonomy, particularly for bacteria, is largely defined at the genetic level. Yet the information encoded in genetics propagates through many levels of biological organization: genes determine patterns of gene expression and protein production which, in turn, influence cellular physiology and behavior which, in turn, influence the collective morphology of growing populations. It is unclear how far up this hierarchy that the lower level taxonomic information remains preserved. Here, we show that bacterial genera can be identified from differences in the three-dimensional topographies of their growing populations. Using coherence-scanning white-light interferometry (WLI), which is capable of nanometer-resolution, we measure bacterial population topographies of 77 different clinical isolates representing four different genera. We then extract ten biophysically relevant features describing their surface structure. Using a simple machine-learning classifier, these topographic features identified bacterial genus with 97% accuracy, demonstrating that genus-level information remains present at the scale of population morphology. We hypothesize that these topographic “fingerprints” arise because differences in the underlying biophysics of bacterial growth propagate upward into measurable differences in population structure. To test this hypothesis, we performed biophysical simulations of bacterial population growth and applied our experimentally trained classifier directly to the resulting simulated topographies. By systematically modifying simulation parameters, we determined which changes were necessary to generate topographies classified as each of the experimentally observed genera. We find that a relatively small number of differences in cellular growth, cell-surface interactions, and initial conditions are sufficient to recreate the genus-associated differences observed experimentally, and we then show that using the topographic features from the simulations we can recover the underlying parameters used to simulate the topographies. Together, these results show that differences in lower-level bacterial behavior can propagate into distinct population-scale morphologies, and that taxonomic information can remain recoverable far above the molecular scale at which bacterial identity is conventionally defined.

## 1 Introduction

Bacteria have colonized Earth. They are present in nearly every environment, including many that are hostile to life. It is estimated that there may be millions or billions of bacterial species and hundreds of thousands of bacterial genera [1]. These estimates come from genetic studies; while bacteria can be classified into large groups based on cell morphologies, e.g., rod-shaped, spherical cocci, comma-shaped, among others, classification based on cellular morphology, long the sole approach of taxonomy in the life sciences, has proven challenging. Further, while morphological differences in group formation exist, such as the growth of chains of cells versus clusters of cells, classification beyond which mechanism is present is difficult [2, 3]. However, this observation conceals two facts. First, the current best estimates suggest that bacteria spend most of their time in biofilms [4], suggesting that the morphological characteristics of multicellular bacterial populations may be more relevant than those of single cells. Second, and in a related vein, identifying animal genera on the basis of individual cell morphologies would be a highly non-trivial task as well. Further, it is known that colonies can provide some clues as to the underlying taxonomy, from the colony color and morphology. However, at first glance, colonies appear to lack sufficiently unique morphological characteristics necessary to categorize bacteria on their basis. For example, while distantly related species may appear visually different, closely related species or genera may be expected to produce visually similar colonies.

However, it is worth questioning if this is in fact true due to the hierarchical nature of organisms. Different bacterial species can be identified on the basis of a sufficient amount of shared DNA. That DNA determines what proteins are made, as well as the relative frequency of each protein. The proteins made and their abundances determine cell physiology and behavior. Finally, cell physiology and behavior, along with their biophysical effects, determine the multicellular structure of a colony or biofilm. In principle, as every level proceeds based on the details of the level below it, every stage of the entire hierarchy is connected back to the genetic level, potentially enabling higher levels to distinguish lower levels. In fact, MALDI-TOF (Matrix-Assisted Laser Desorption/Ionization Time-of-Flight Mass Spectrometry) has proven capable of identifying bacterial genus with high accuracy on the basis of identifying proteins and their abundances [5].

While this fact shows that genera are distinguishable at least one step up this hierarchy, it is unclear if subsequent steps will behave similarly. While higher levels depend on the details on the lower levels, emergent phenomena can make higher levels distinct from lower levels. Phenotypic plasticity and stochastic gene expression are two such examples; the same genotype can produce different phenotypes as environmental conditions, cellular history, and intrinsic fluctuations introduce information not contained in the genetic sequence alone. As information from these additional sources accumulates at higher levels of organization, distinctions present at lower levels may become increasingly obscured. Similarly, it is difficult to predict the details of a colony by studying single cells in isolation as their multicellular interactions play a fundamentally important role. As a result, two broad possibilities may prevent colony morphologies from identifying bacterial genera. First, colony growth may be too chaotic. The initial conditions or the subsequent growth of bacteria may be too stochastic and as a result there may be as much variation within a genus as across genera. Second, there may be emergent forces that erase detailed information about the bacteria that produce a colony. For example, colonies are well known to be viscoelastic; viscous relaxation could erase distinguishing details from colonies, leaving them as simple indistinguishable films. Thus, it remains unclear if colonies possess enough distinct features to connect their morphologies to the genera of the bacteria that produced them.

Fortunately, recent work has demonstrated that the underlying biophysics of bacterial growth produces bacterial population topographies that are rich with information about the bacteria that create them. Measurements of the topography of a bacterial population have been used to understand the biophysics of the vertical growth of a colony [6, 7], as well as the biophysical connection between horizontal and vertical growth [8, 9]. As many recent works have demonstrated the link between bacterial biophysics and population morphology [10–17], we hypothesized that different underlying biophysics (e.g., differences in vertical growth rates, or cell-surface friction) will manifest topographic “fingerprints” that can be connected to the ID of the bacteria that produced them. In other words, differences in bacterial behavior ought to be reflected in bacterial population topographies; so long as bacteria of the same taxonomic group behave similarly enough, their topographies should show similar signatures. This raises two natural questions of resolution. First, how genetically similar can two bacteria be while still producing distinguishable topographies? And second, how different can the topographies from two different strains of the same species be? As a first proof of concept, we begin at the genus level, asking whether topographic signatures are sufficiently conserved within genera while remaining distinct across genera.

We sought to determine if differences in topographies are robust across genera, thus enabling bacteria to be identified through their topographic morphologies. To test this, we measured the surface topography of bacterial populations using coherence-scanning white-light interferometry (WLI). This technique has been shown to be capable of detecting cell growth[18–20], differentiating populations with and without microbial warfare[21], and revealing key insights into the dynamics of bacterial colony growth[6–9, 22]. For the identification task itself, we chose interpretable features and a simple machine learning (ML) classifier rather than deep learning techniques to ensure that classification inputs and outputs remained mechanistically understandable, enabling us to relate those inputs to downstream biophysical characteristics while minimizing the risk of learning spurious dataset-specific signals (e.g., background noise) that may not generalize. Thus, we next identified a host of topographic features that are directly relatable to the biophysics of vertical and horizontal bacterial growth. We then used a support vector classifier to validate our hypothesis that topographies contain rich identification information. This method for identifying bacterial genera (ID) from topographic features obtained 97% accuracy in genera after 4 hours of incubation, validated across 77 clinical isolates consisting of four gram-negative ESKAPEE (highly antibiotic resistant group of pathogens commonly responsible for nosocomial infections) species. To understand why high-resolution topographic analysis is as effective as it is so quickly and with only one single measurement, even for conventionally hard classifications like closely related genera, we performed biophysical simulations of colony growth. We then applied our experimentally-trained ML classifier to the biophysically simulated topographies to determine which simulation parameters are necessary to produce topographies classified as different genera. We found that a small number of growth parameters and initial conditions, representing biological and physical differences between isolates, were capable of recreating the differences observed in experimental topographies.

## 2 Results

### 2.1 Isolates measured

We selected four genera to test, with one species from each: *Acinetobacter baumonii*, *Pseudomonas aeruginosa*, *Escherichia coli*, and *Klebsiella pneumoniae*. This set of four gram negative bacteria are all rod-shaped and include a pairing of two very closely related genera *Escherichia coli* and *Klebsiella pneumoniae*. We worked exclusively with clinical isolates to avoid the potential uniqueness of lab strains that have been grown repeatedly in rich media.

To measure bacterial population topographies, we inoculate 5 µl of liquid-suspended, density-standardized clinical isolates onto agar plates. We then incubate for four hours and measure the population topography through a single, rapid (*∼*3 seconds) scan with a Zygo NewView WLI.

### 2.2 Extraction of biophysical features for machine learning

As we have hypothesized that differences in the underlying biophysics of bacterial growth will produce fingerprints of different bacterial genera, we specifically measured features of the topographies that can be directly tied to the underlying biophysics of bacterial growth on a surface.

To understand these features, we first must identify the three distinct spatial regions found in each topography (Fig. 1). The first is outside the population, i.e., where there are no bacteria. This is the agar on which the bacterial population rests. Identifying this region is essential for measuring absolute sizes of features, as well as for correcting any background tilt and agar substrate deformations. The second region is where the highest density of cells is found: the coffee ring, named for the ‘coffee ring effect’. The coffee ring effect[23, 24] is a physical phenomenon resulting from a drying drop containing colloidal particles with “pinned” edges; surface tension drives fluid flows from the drop center to the drop edges, carrying anything suspended in the drop, including bacteria, to the pinned edges. After the drop dries, bacteria are densely packed in the coffee ring. The third region is the area inside the coffee ring called the homeland. For the cell densities that we inoculate, the homeland does not start with a complete cellular monolayer, but instead is initially sparse. The differences in cell density create different dynamics[25] that can impact the topographies they produce.

**Fig. 1.**
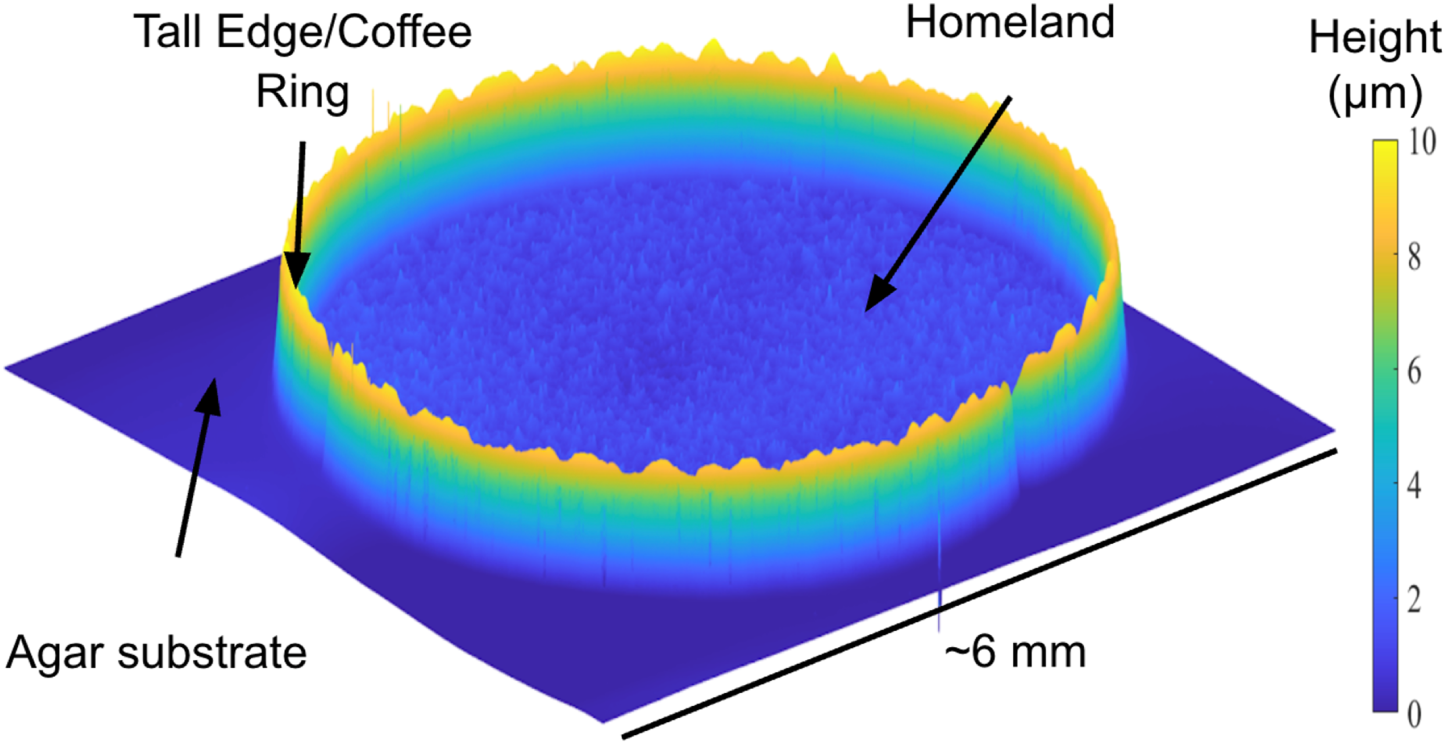
Bacterial population topography measured using white light interferometry (WLI). An *Escherichia coli* clinical isolate was deposited on an agar pad in a 5 µl drop. Population topography was measured using WLI and is shown, with distinct regions such as the coffee ring, homeland, and agar substrate (devoid of bacteria) marked.

We identified features that we expected to relate to the biophysics of bacterial population growth. For example, the colony contact angle was shown to have a strong correlation with horizontal growth [8], and vertical growth was shown to depend strongly on a small number of parameters [6, 7]. Features come from both the coffee ring and the homeland (details on all of the features and their calculation can be found in the Methods section).

### 2.3 Simple Approach to Machine Learning

To identify genera from the features extracted from a population topography, we use a simple ML classification method, a linear-kernel support vector machine (SVM). The only alteration of the features is a z-scoring to increase the speed of convergence[26]. While we utilize a relatively large biological sample set for an initial study with nearly 20 isolates of each genus (Fig. 2), we are still considered to be data-limited by ML standards. Thus, to maximize our training and testing sets, we use the leave-one-out cross validation (LOOCV) technique to provide the reported accuracy for each sample. We perform two replicate experiments for every strain. The prediction is the species with the largest model confidence score across the two replicates. We also perform a two-stage approach, to help improve our “image to number-of-classes” balance. The first stage groups Enterobacterales together (*E. coli* and *K. pneumoniae*) and the others together (*A. baumanii* and *P. aeruginosa*) then the second stage breaks each group into its two respective species. See Methods for more info.

**Fig. 2.**
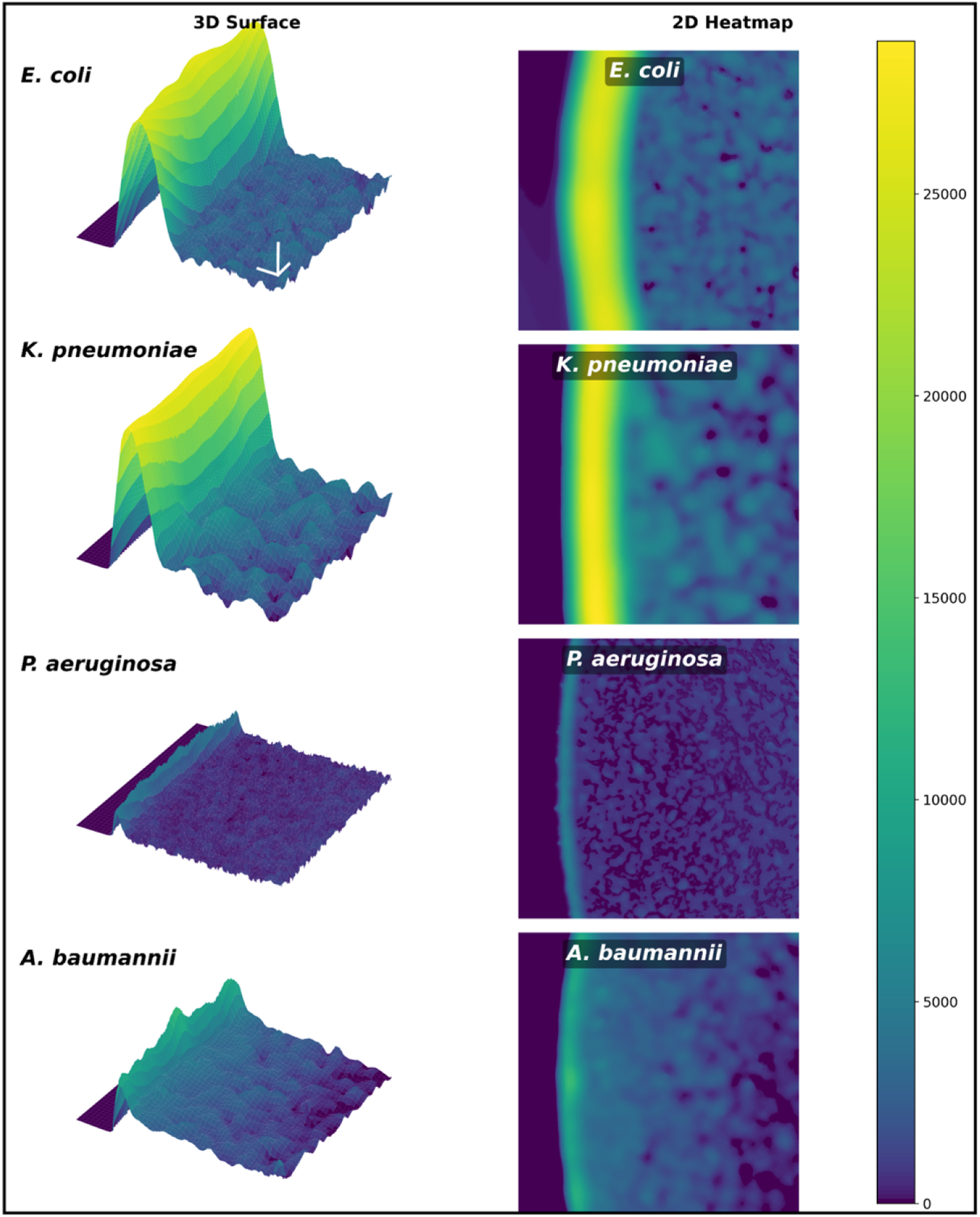
Representative images of four species tested including *Escherichia coli*, *Klebsiella pneumoniae*, *Pseudomonas aeruginosa*, and *Acinetobacter baumannii* (top to bottom, respectively). A full 3D image and 2D heatmap are shown for each species. The color bar on the far right defines the heights for all images reported in nanometers. 3D scale bars are show in the *E. coli* 3D surfaces; the X- and Y-direction bars represent 40 µm and the Z-direction bar represents 6 µm.

### 2.4 Taxa Identified with Ultra High Accuracy and Short Incubation

Using the approach described above, we correctly sorted 97.4% of strains into their correct taxonomies (75/77 - Fig. 3). The only two errors came from a strain of *Pseudomonas* that was incorrectly called *Escherichia* / *Klebsiella* in the first ML stage, and an *Escherichia* strain that was incorrectly called *Klebsiella* in the second ML stage.

**Fig. 3.**
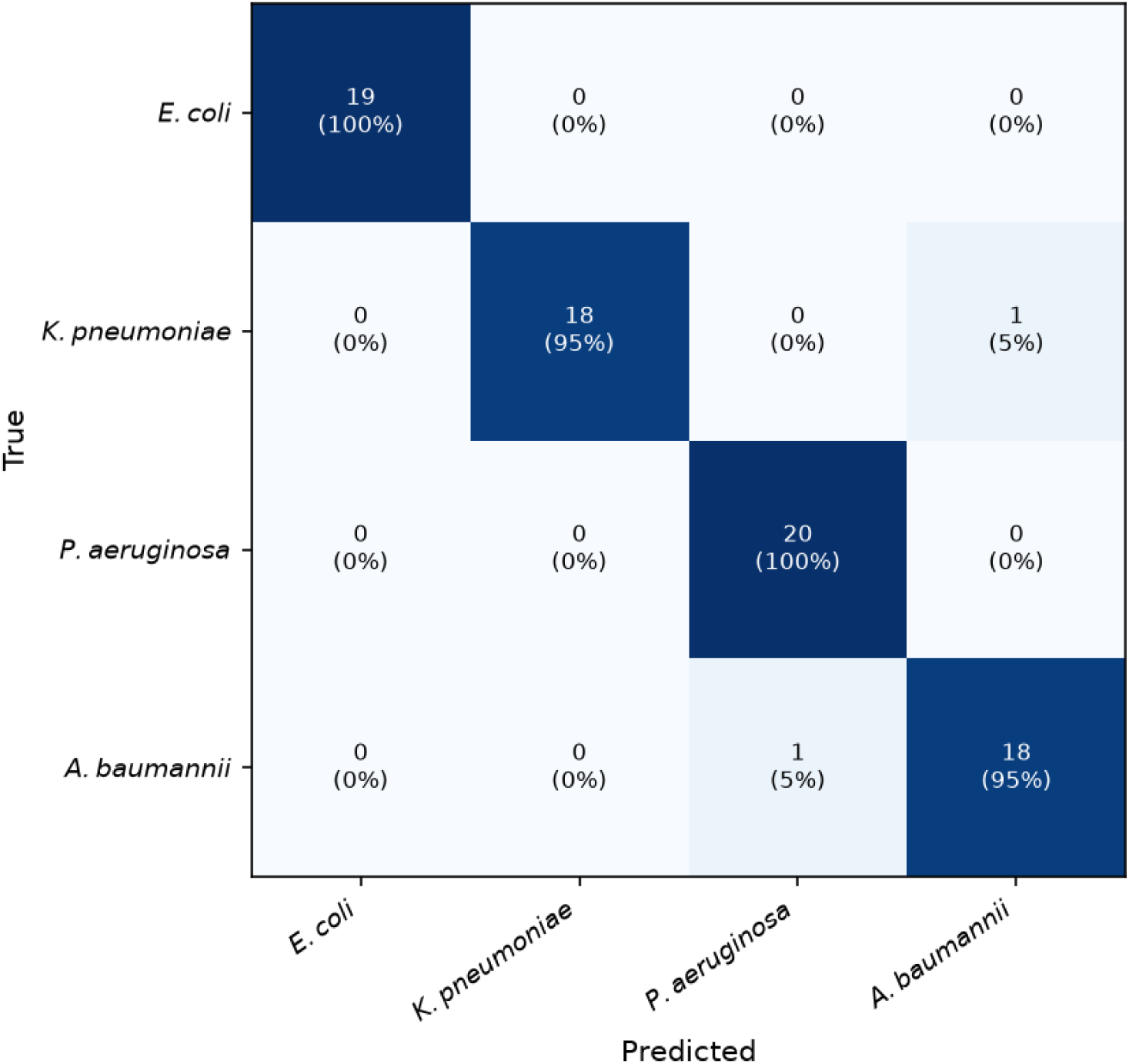
Confusion matrix showing predicted and true genus, and the percentage of each call.

#### 2.4.1 A consistent set of features accurately separates taxa

We next sought to minimize the information needed to classify the samples so that our later investigation into these features is focused. Therefore, we performed RFE (recursive feature elimination) and varied the number of features included from 1-10 (Fig. S6) out of 10 proposed. We find that accuracy is optimized if six features are kept in the first stage; the second stage accuracy is optimized with six features for distinguishing *Escherichia* and *Klebsiella*, and two for distinguishing *Acinetobacter* and *Pseudomonas*. For fewer than four features, the accuracy of each stage falls (for one parameter, the accuracies are 81%, 95%, and 89% for stage 1, *Acinetobacter* vs. *Pseudomonas* and *Escherichia* vs. *Klebsiella*, respectively). Thus, we focus our analysis here on cases where the optimal number of features are included in the analysis.

There is only one feature that appears in every classifier: the contact angle. The contact angle is a rich, emergent property that contains information about how cells interact with each other and the surface, and it is apparently useful for distinguishing genera. Along with the contact angle, the stage one classifier is optimized by the coffee ring width, variance in homeland height, coffee ring height, coffee ring height standard deviation, and the produce of the coffee ring height multiplied by the homeland height. The *Acinetobacter* vs. *Pseudomonas* classifier is optimized with the contact angle and coffee ring Hurst exponent. The *Escherichia* vs. *Klebsiella* classifier is optimized with the coffee ring width, the coffee ring height, the coffee ring Hurst exponent (a metric that characterizes if fluctuations are random walk-like or not; see methods for more details), the contact angle, and the ratio of the coffee ring height to the homeland height. It is unsurprising that the first stage features the coffee ring height given that this height is a function of both the complex initial conditions of the bacterial population (which itself is a function of many parameters including cell-cell adhesion, cell shape for packing, cell-surface adhesion, and more) and non-linear fluid dynamics during growth, parameterized by metabolism, cell death, and nutrient diffusion through the extracellular matrix [7, 24]. For distinguishing *Escherichia* and *Klebsiella*, all features come from the coffee ring; for distinguishing *Acinetobacter* and *Pseudomonas*, all features except one come from the coffee ring.

Further, we attempted to perform the two stage ML approach but this time with only one feature (while iterating through all features). Remarkably, the ratio of the coffee ring height to the homeland height produces an accuracy of 79.2% on its own, demonstrating that a simple metric containing information about the coffee ring and homeland is highly capable of distinguishing genera (Fig. S4).

Thus, while we have shown that topographic fingerprints can distinguish the genera of the bacteria that produced them, we next seek to understand why.

### 2.5 Importance of High Resolution to Read Biophysical Information

To understand why bacterial population topographies contain information about the genera that produced them, we first sought to understand which aspects of the topographies contribute the most to these features. To investigate, we re-trained our models on subsets of the features initially used. In particular, we evaluated four different subsets of features: coffee ring features only, homeland features only, lateral features only, and vertical features only. Lateral features refers to features that do not explicitly include the vertical height, such as the coffee ring width or power spectrum values. Vertical features include only features calculated from heights alone; thus we did not include contact angle or the Hurst exponent, which include vertical and lateral information. We then repeated the LOOCV-RFE approach detailed above, producing accuracies for these different conditions.

We found that two modifications had minor impacts on our accuracy, and two had major impacts. Using the coffee ring only or vertical features only decreased our overall accuracy to 93.5% and 90.9%, respectively. Conversely, we found that using features from the homeland only or using lateral features only had a catastrophic impact on accuracies, dropping them to 59.7% and 54.5%, respectively.

Overall, these results suggest that vertical information plays the dominant role in the accuracy of WLI in identifying bacterial genera. This is not surprising as WLI is capable of super-resolution in the vertical direction, but is diffraction-limited (and more relevantly, limited by the NA, numerical aperture, of the interferometer objectives) in the lateral direction. As a result, it is likely that vertical features carry more information than horizontal features for WLI. Nonetheless, these results show that, in fact, topographies can be linked to the genera of the bacteria that produce them, so long as the topographies are measured with sufficient resolution.

Why are vertical features so distinguishable? To start, it is clear that the population’s topography, and thus its vertical features, emerges from the population’s initial conditions and subsequent growth. For example, the coffee ring height will depend strongly on the vertical growth rate [6, 7] and the initial height of the coffee ring, while the contact angle depends strongly on the competition between vertical and horizontal growth rates [8]. Thus, the fact that vertical features are so distinguishable implies that isolates within each genera produce similar initial conditions and grow similarly, and that these initial conditions and subsequent growth differ across genera. We next seek to understand if initial conditions vary across species.

#### 2.5.1 Initial conditions vary by genera

The initial condition arises after the inoculation drop dries. Bacterial suspensions are known to exhibit the coffee ring effect, i.e., the phenomenon in which suspended matter in a drying drop is pushed to the drop’s edge during drying. The coffee ring effect is a well-studied phenomenon, and is known to be affected by a wide range of phenomena. However, rather than attempting to understand how these deposits form in detail, our goal is to determine if different genera have different coffee ring initial conditions, if all genera have the same initial conditions, or if initial conditions vary substantially without correlation to genera.

Indeed, we measured the coffee ring immediately after drying (*∼*5 minutes after inoculation) across 75 samples. We found (Fig. S1)that the coffee ring height varies little across genera (*<* 1 µm), while the coffee ring width varies substantially (*∼* 35 µm, albeit with large standard deviations; see Fig. S1). Thus, some of the distinguishable features could arise from differences in initial conditions which themselves stem from more complicated properties of the underlying bio-physics.

#### 2.5.2 Biophysical Simulations Produce Realistic Topographies

To move beyond features and determine which broad biophysical parameters most influence these topographies, we perform group level, simple bio-fluid simulations. Vertical and horizontal growth are well-studied [6–8, 10, 11, 17, 27, 28], and recent works have shown experimental results are very accurately reproduced with simple mathematical models [6–8]. Thus, if our hypothesis is correct, simulations of bacterial population growth from an inoculation-like initial condition ought to produce the sorts of topographies observed in experiments. Further, changes to the simulated growth parameters should lead to changes in topographic features, so the growth parameters themselves ought be broadly recoverable from the topographic features.

To test these hypotheses, we performed biophysical simulations of growing populations. Three-dimensional simulations of the macroscopic growth and morphology of bacterial colonies were inspired by [6–8]. The simulations initialize a 3000 by 3000 µm spatial grid with a central 1 mm radius coffee ring of biomass and a designated number of smaller internal biomass clusters seeded randomly within the ring. The height of the colony at any point on the grid evolves over time (simulating 4 hours of growth) according to the following equation:

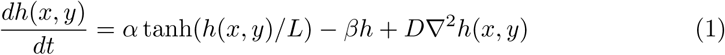

where *h*(*x, y*) is the height of the population at position (*x, y*), *α* is the vertical growth rate, *β* is a death or decay rate, *L* is the maximum height to which exponential growth is possible (due to the competition between nutrient uptake and diffusion of those nutrients), and *D* is the coefficient that governs the spread of bacteria across the surface. This expression was validated to replicate experimental data in [7] and [8]; values of *α*, *β*, *L*, and *D* are taken from previously extracted experimental values. Finally, the initial conditions were varied as well: the initial height of the coffee ring of cells, the amount of biomass inside the coffee ring, the number of clusters, and the initial width of the coffee ring were all varied. The exact values of the simulation parameters for our tested species is both unknown and immensely time-intensive to measure. Thus, simulations were performed across *∼*30 thousand combinations of these parameters to generate a broad library of theoretical colony topographies, which are output as 2D height arrays (Fig. 4).

**Fig. 4.**
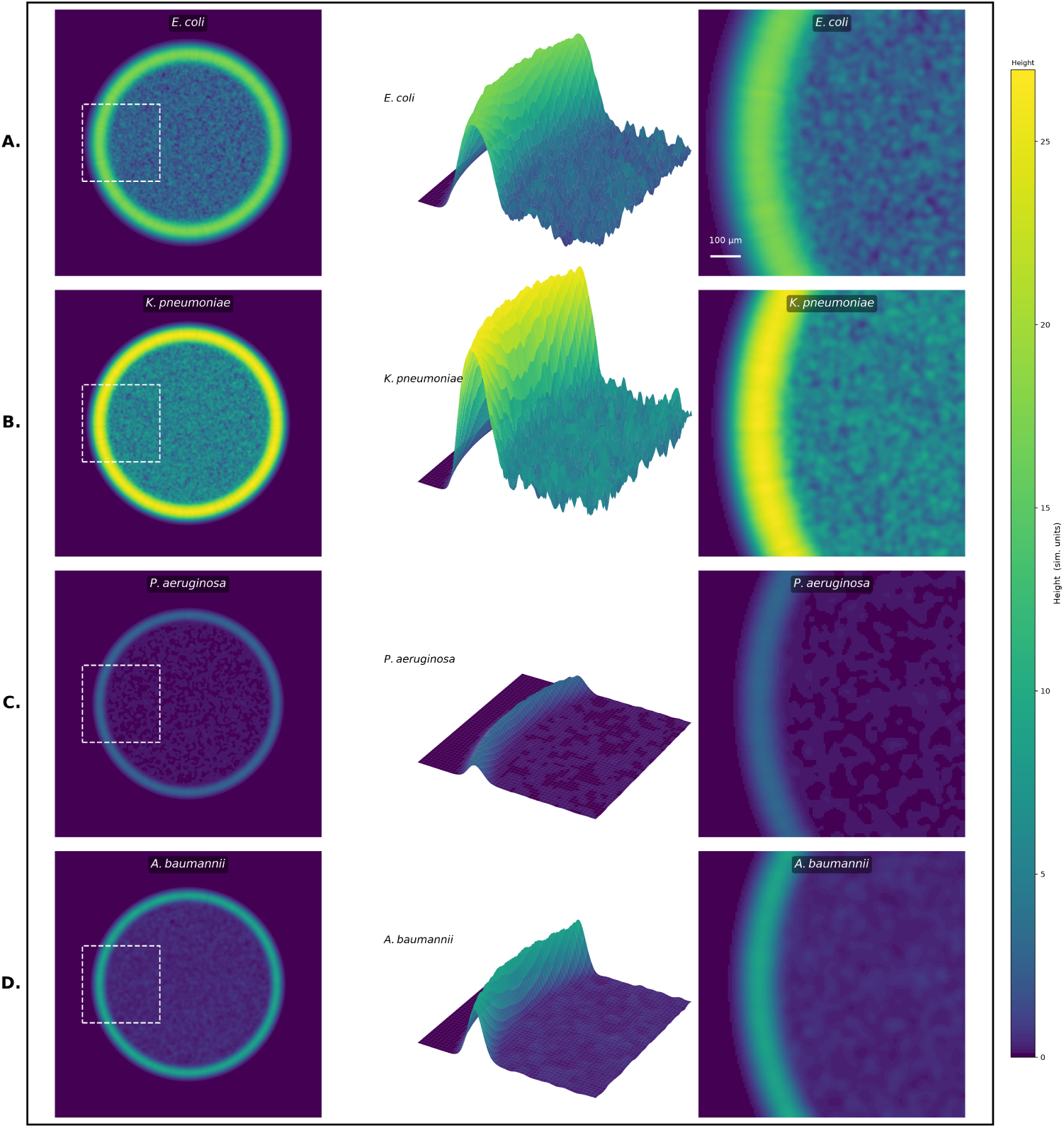
Visualizations of biophysical simulations of *Acinetobacter*, *Pseudomonas*, *Klebsiella*, and *Escherichia* bottom row. Compare visually with Fig. 2

We first sought to determine how the simulation parameters affect the biophysical features. To do so, we performed a one-way analysis of variance (ANOVA) and calculated eta-squared (*η*^2^) effect sizes, which measures how much of the variance in a given feature is explained by a particular simulation parameter (Fig. S2). We find that some simulation parameters affect many features (e.g., *α*, which has *η*^2^ *>* 0.2 for five different features, and *D*, which has *η*^2^ *>* 0.5 for two features), while others have little impact (in particular: L, the initial coffee ring height, and the number of homeland clusters all have *η*^2^ *≤* 0.12 for all features).

The same features that were used to identify experimental IDs were then fed into the experimental classifier, producing predicted species for each topography. The classifier sorted topographies into all four genera. While the values of the features were not always identical to those from experiments, plots of the simulated values of these features versus the experimental values demonstrate that the rank order is largely preserved (Fig. S3).

To understand why simulations were classified as different genera, we compared the simulation parameters that produced them (e.g., we compared the *α* values for *Klebsiella* simulations to those of *Escherichia*, *Psuedomonas*, and *Acinetobacter*). We analyzed all comparisons between genera and parameters (42 in total; six genera-to-genera comparisons for seven parameters), and performed t-tests. We found that almost every parameter comparison was highly statistically significant (*p <* 0.001, Welch’s t-test; see Fig. S5 for all comparisons). In fact, only five comparisons were not significant (*P >* 0.05). Further, while most of these differences are small (*<* 10%), many were large. Of the 42 total comparisons, 20 differ by more than 10% and, of those, 16 differ by more than 30%. All combinations of species had at least two parameters that were highly statistically significantly different that differed by at least 10%.

A one-way ANOVA with *η*^2^ effect sizes was used to identify which simulation parameters varied most strongly across the four classifier-assigned genera in the pure simulations. All simulation parameters reached statistical significance (*p <* 0.001). The growth rate, *α*, was the most dominant driver of genus assignment (*η*^2^=0.24), followed by the diffusion coefficient, *D* (*η*^2^ = 0.10), the fraction of biomass initially in the coffee ring (*η*^2^ = 0.08), and the initial width of the coffee ring (*η*^2^ = 0.07). *β* and *L* contributed the least (*η*^2^=0.019 and 0.015, respectively); these results were expected, as *h < L* at most locations during these four hours of growth, preventing either of these parameters from heavily influencing the results.

Further, four natural groupings of simulation parameters emerge. *Pseudomonas* has the lowest average growth rate (*p <* 0.001). 77% of simulations classified as *Pseudomonas* had *α ≤* 0.6, while only 21% of our simulations overall had growth rates that low. *Klebsiella* has the highest average growth rate (*p <* 0.001); all *Klebsiella* classified simulations had *α ≥* 1.6 even though only 10% of our simulations had growth rates that high. *Acinetobacter* has the largest initial coffee ring width (*p <* 0.001); 97% of simulations with an initial coffee ring width of 120 µm were classified as *Acinetobacter*. These results suggest that for the four genera studied, the variance of biological properties within a single genus is smaller than the variance across genera. As the topography depends on all of these parameters, higher level topographic features that emerge from these simulated cell-level properties are able to distinguish genera. However, at this stage we have only looked at classification into broad categories (genera), and while the resulting topographies from the simulations appear similar to the matching genera by eye, simulations do not have ground truth labels for genera. However, if the lower level information is propagated up through the topographic features, we ought to be able to determine simulation parameters directly from topographic features.

#### 2.5.3 Biological information encoding

Finally, we sought to directly test if topographic features can be linked to the lower level traits of the simulated bacteria that produced them by directly predicting simulation parameters from extracted features. A neural network classifier was trained to predict seven simulation parameters (*α*, *β*, L, D, fraction of cells initially in the coffee ring, number of clusters initially in the homeland, and the initial width of the coffee ring) from nine topographic features (see methods for more details).

We find that we can predict the exact class of simulation values with high accuracy. Five of the seven parameters (*α*, *D*, the initial fraction of cells in the coffee ring, the initial number of clusters in the homeland, and the initial width of the coffee ring) are predicted correctly (i.e., the exact value is predicted) over 97% of the time. *β* is predicted correctly 95% of the time, and L is predicted correctly 89% of the time; this makes sense as *β* and L hold little influence on topographic features over the time scales we investigated. Overall, these results suggest that the reason topographic features can positively identify genera is because they are closely linked to cellular behavior.

## 3 Discussion

Here we have shown, using white-light interferometry, that bacterial surface topographies are sufficiently unique to distinguish between four gram-negative rod-shaped species with very high accuracy. The results here demonstrate that interferometry can differentiate between these similar bacteria due to its ability to perform high resolution vertical measurements. The topographic features that proved most useful were typically those directly connected to the underlying biophysics, such as the contact angle or the height of the coffee ring. Further, biophysical simulations demonstrate that consistent differences in initial conditions and growth parameters produce topographies that can be classified distinctly as all four genera studied. Thus, the biophysics of bacterial growth—how bacteria push each other laterally and vertically as they grow on agar—produces biophysical fingerprints in bacterial population topographies, capable of distinguishing or categorizing bacteria on their basis.

Of course, thousands of bacterial genera, and tens of thousands of bacterial species, have been identified. It remains to be seen if genera could be positively identified if the scope were increased to tens, hundreds, or even thousands of genera. However, there are some reasons for optimism. First, all four species studied here are rod-shaped bacteria that are *∼*1.5 µm in length. We expect that different shapes of cells (e.g., *Streptococcus* which are coccoid) and different sizes of cells (e.g., *Mycoplasma genitalium*) could be easy to distinguish. Further, while we focused on a single media and a single time point, more genera and species could likely be distinguished by growing on multiple media or for different lengths of time.

Focusing towards human health, bacterial infections are the second most common cause of death worldwide [29]. Treating these infections effectively requires knowledge of the bacterial identity, both for pathogensis information and for setting *in vitro* concentration standards for antimicrobial testing. Our foundational principle demonstrated here is that biological differences between different species determine their physical and chemical characteristics that are responsible for how they interact with each other and their surroundings and how they grow spatiotemporally. Fortunately, out of the tens of thousands of bacteria, there are only 1500 that are pathogenic to humans [30], and only a small fraction of those that are responsible for most infections. For example, one study found that the “ESKAPE” pathogens, including three of the four we tested, were responsible for *>* 70% of infections in one hospital’s emergency room [31]. Thus, while the work presented must be tested against more than four genera, it need not be capable of distinguishing any and all possible comparisons to prove useful.

Finally, while we note that these topographic fingerprints emerge in well-controlled laboratory experiments, in nature, most biofilms are polymicrobial, which would preclude this sort of analysis. Nonetheless, future work may try to determine if monoclonal biofilms grown in nature also produce unique fingerprints, or if these fingerprints emerge from the unnatural laboratory conditions, e.g., a very large number of cells completely isolated from other species and held at an ideal temperature.

## 4 Data and Code Availability

Data and code will be available at https://github.com/peter-yunker.

## 5 Acknowledgments

This work was, in part, funded by the Department of Veterans Affairs (VA) grant BX005985 to DSW and PJY. PJY was also funded by a grant from the National Institutes of Health, National Institute of General Medical Sciences (grant no. 1R35GM138354-01). DSW was also funded by a Burroughs Wellcome Fund Investigator in the Pathogenesis of Infectious Disease award.

## 6 Methods

### 6.1 Strains

Clinical isolates of the indicated species were acquired from the Investigational Clinical Microbiology Core of the Emory Antibiotic Resistance Center. These are unique patient isolates, i.e., each strain was the only strain isolated from a single patient.

### 6.2 Inoculation on agar pad

1.5% Mueller-Hinton agar (MHA) plates were prepared with the Clinical & Laboratory Standards Institute (CLSI) resistant breakpoint concentration of antibiotic for each bug-drug combo (see Table 1) and without antibiotics for pathogen identification testing. For the best metrology measurements, we avoided large-scale (1s-10s of microns out-of-plane, millimeters to centimeters laterally) ridges which can significantly impact measurement and may arise due to differential contraction rates with spatial cooling heterogeneities.

**Table 1.**
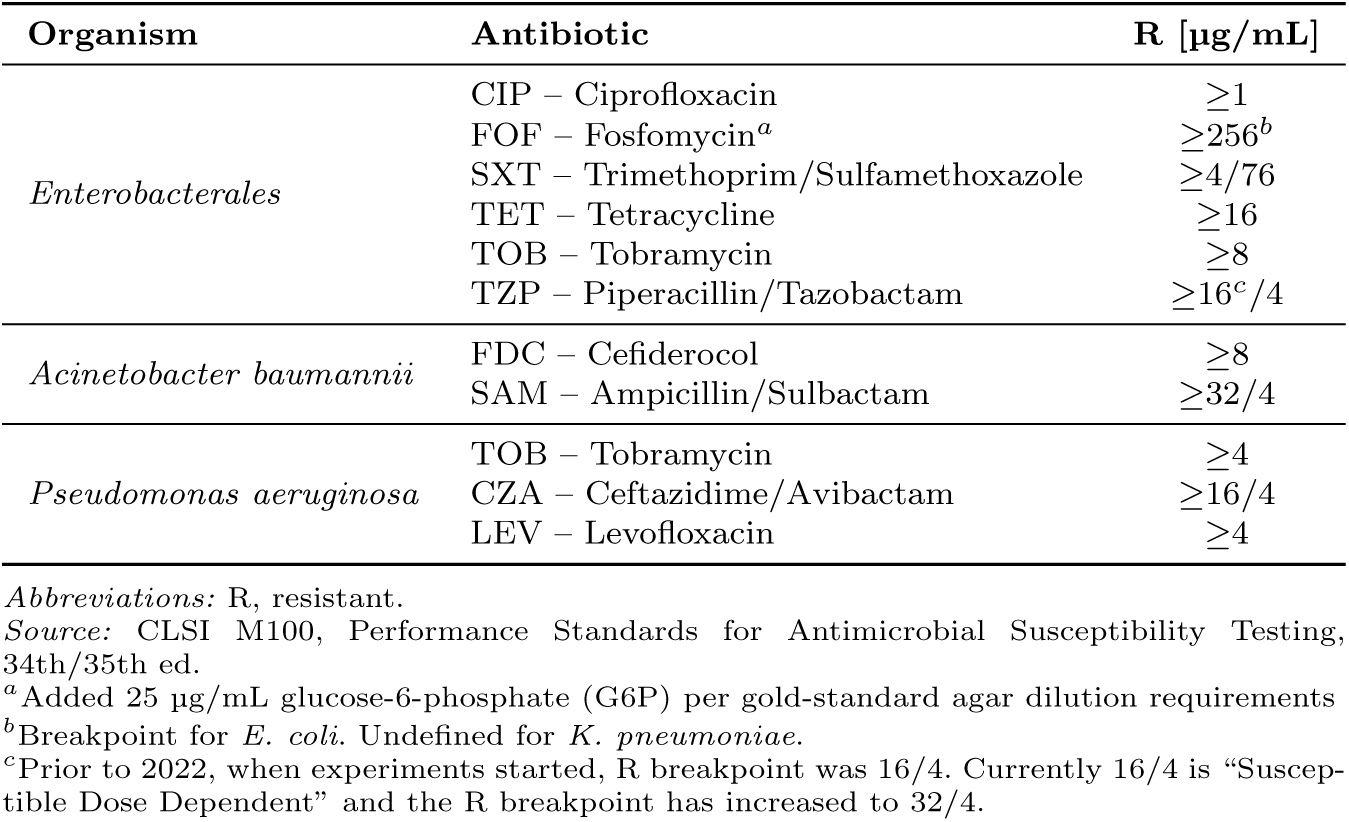
CLSI M100 MIC Resistance Breakpoints (µg/mL) by Organism and Antibiotic.

For antibiotic susceptibility testing, isolates were streaked onto MHA plates and incubated overnight. Five colonies were selected the next day and mixed with 100*µl* of phosphate buffer solution (PBS) in a 96 well plate. The *OD*_600_ was measured in a well-plate reader and the solution diluted to an OD of 0.05. This yields approximately 200,000 cells/*µl*.

For pathogen identification, the *OD*_600_ was measured using 1 mL cuvettes in a portable spectrophotometer with a MHB background. The overnight was diluted to an *OD*_600_ of 0.1 which yields approximately 100,000 cells/*µl*.

5 *µl* were inoculated as a droplet onto the MHA with special care taken to not poke the agar with the pipette tip. The spot was left to dry for 5-15 minutes and then taken for measurement.

### 6.3 Topography measurement

For pathogen identification, we used a Zygo ZeGage Pro interferometer a 10X Mirau interferometry objective (NA=0.30, 0.87µm/px).

Three distinct regions of topography exist once the inoculated droplet dries: the agar substrate, the “coffee-ring”, and the “homeland”. The agar substrate provides a reference point to calculate absolute heights of the topography created by the microbes. The coffee-ring effect deposits cells in a dense pile along the edge of the pinned droplet and thus defines the edge of the inoculated sample. The homeland is the region contained within the coffee-ring/colony edge where cells are sparse at the relevant cell densities. We ensure that data is collected from each region. With translation to a clinical device in mind, we minimize measurement time to 3-5 seconds by using objectives that achieve a sufficient total field of view without stitching images. Simultaneously, we took care to consider the necessity of single cell information (higher lateral resolution) and the specular reflection limit (related to numerical aperture - ‘slope limit’) required of meaningful data on the scales deemed important for the classification. In general, higher magnification objectives provide higher lateral resolution and an increased slope limit at the cost of the total field of view and thus measurement time. Each objective is capable of achieving single-nanometer axial (out-of-sample-plane) resolution.

### 6.4 Topographic data analysis

Topographies are cleaned and converted to relative height values instead of the global heights provided relative to the inertial frame of the interferometer. With a meaningful topography, a series of biophysically relevant features were extracted.

Each topography was aligned such that the field of view placed agar on only one side of the image and the coffee ring aligned approximately vertically. Each sample was tilted to ‘null’ the fringes, as is suggested by Zygo. This was done manually on the ZeGage Pro and using the ‘Auto-tilt’ function on the NewView 8000. After collection, some data, mainly along the steep edges of the coffee ring, is missing due to the specular reflection limit. These missing data points are filled by a Zygo Laplacian interpolation algorithm. From these, we removed the global sample tilt and topographic background using an extrapolation of a plane fitted to the agar substrate to obtain heights of the bacteria *relative to the agar*. While the characterization of agar is outside the scope of this manuscript, it is of great concern. Linear approximations of the surface are generally insufficient for fields of view as large or larger than ours, especially without special care taken when creating the substrate.

We separately consider the homeland and the coffee ring *in silico* to calculate the features. The topography is represented as a 2-dimensional array of height values. The coffee ring was consistently aligned with axis 1 (using traditional imaging/array conventions) making it vertical in the array when looking from above. The profilers provided 1000x1000 (ZeGage Pro) and 1024x1024 (NewView 8000) arrays. The coffee ring was aligned such that *∼*150 columns were only agar, available for background fitting. To extract the coffee ring as a quantifiable region, we averaged the three tallest values of each row in the topography array to find the approximate peak of the coffee ring as a function of the vertical axis, collapsing the coffee ring to a 1 dimensional array. For each row, the index of the maximum height was identified along with the nearest indices for heights half the maximum height. The Full-Width Half-Max was calculated to be the difference between the half-max indices and scaled by the pixel spacing. This was averaged over all rows to report a single number. The homeland was taken to be a square region 400x400 pixels along the edge of the array containing the homeland, allowing some separation between the width of the coffee ring and the defined homeland region.

We analyze each relevant region separately, and then make comparisons between the regions. We start with analyzing the homeland using simple metrics built from statistical moments of the height distribution. In particular, we calculate the mean, variance, and coefficient of variation of the topographic heights in the homeland. Then we characterize the same metrics of the coffee ring by first identifying the one-dimensional peak profile of the coffee ring, i.e., around the coffee ring, on its top.

We next further analyze the coffee ring, hoping to take advantage of the large number of cells it contains. Using its peak profile, we analyze how the standard deviation scales in different length bins through the Hurst exponent (Fig. 7), the power law exponent between the measured standard deviations and the bin size. The Hurst exponent measures spatial similarity across length scales. Then we calculate the median full-width-half-max of the coffee ring.

**Fig. 5.**
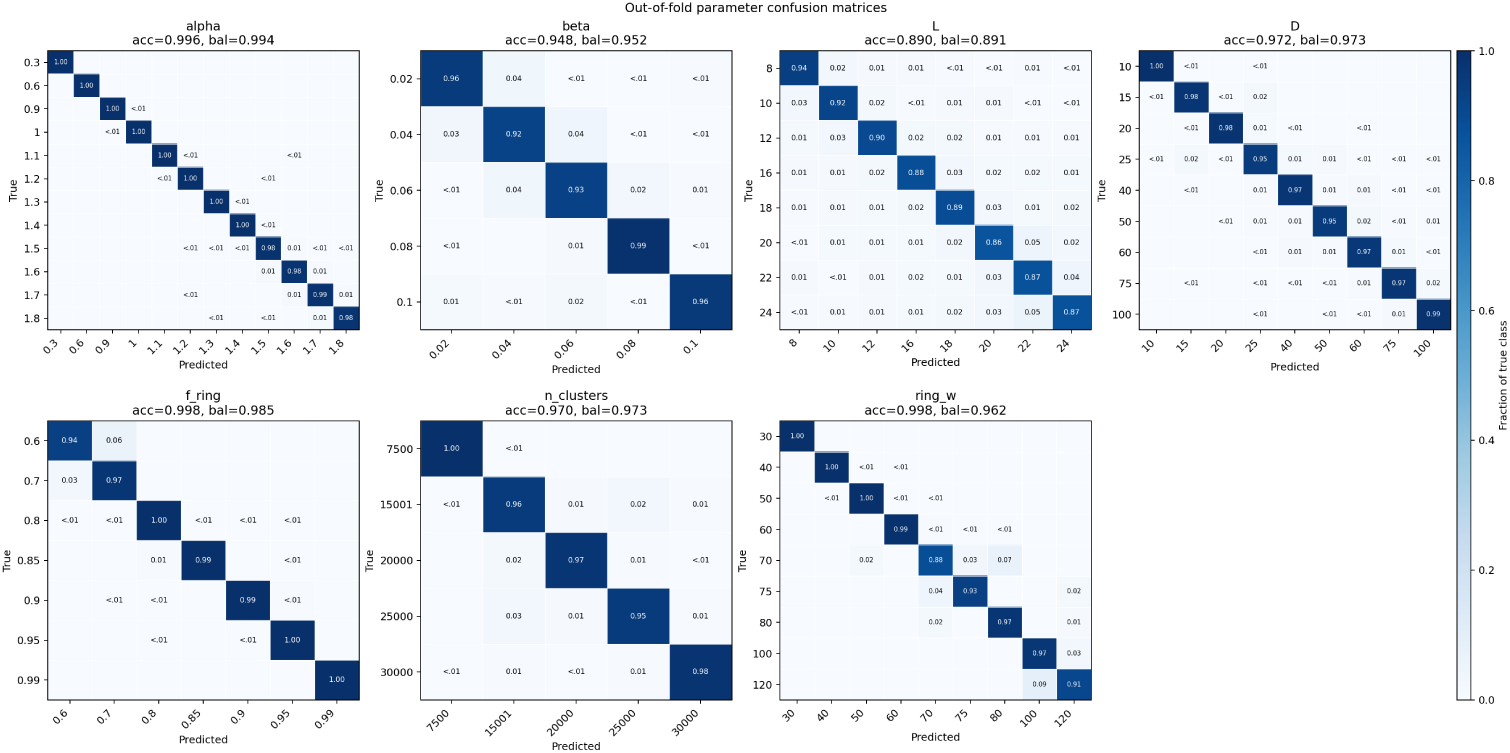
Confusion matrices show the actual and predicted values of simulation parameters.

**Fig. 6.**
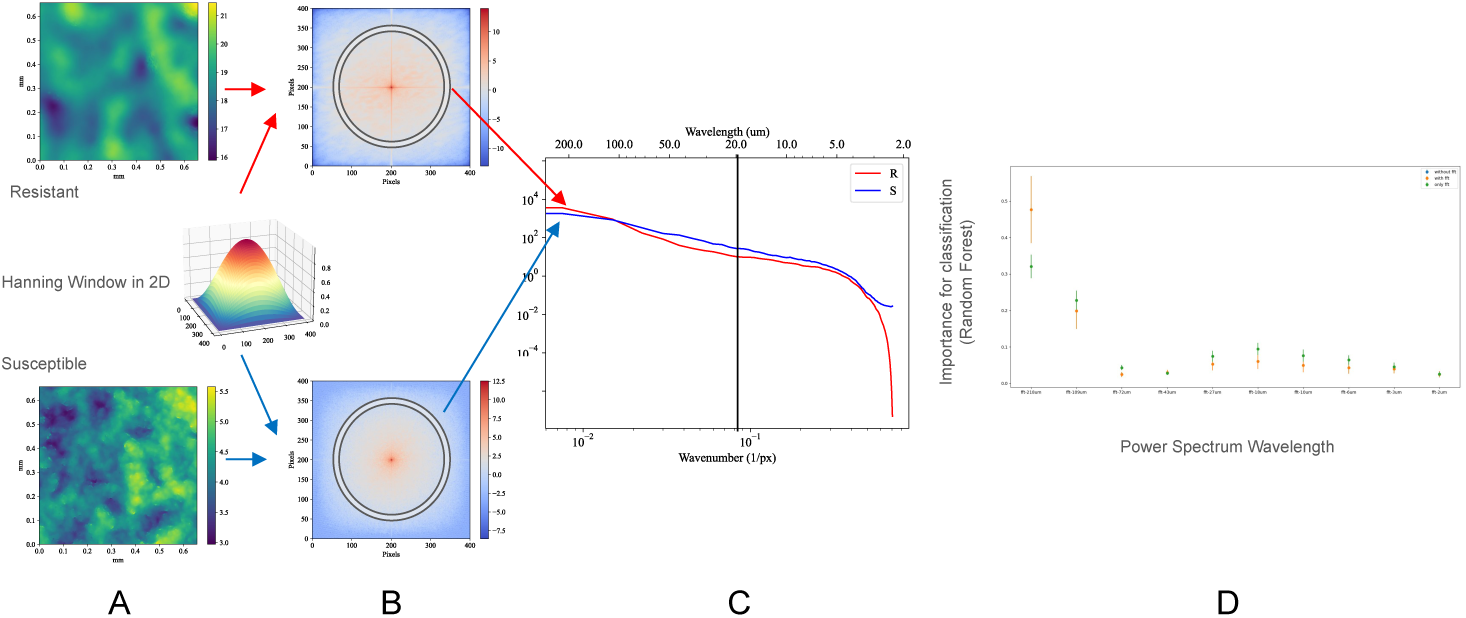
Calculating the power spectrum amplitude. **A**) The homeland for a resistant and susceptible strain are shown. They are multiplied by a 2d Hanning window to account for discontinuities caused by nonperiodicity in the data that results in sharp peaks on the Fourier axes. The logarithm of the complex amplitude of the product is shown in (**B**) along with a representative averaging ring of width dk=3 to calculate the power spectrum in (**C**). **D**) Ten wavelengths sampling the log k-space of the power spectrum were extracted and tested both with the other features and with the only features being the power spectrum amplitudes. For each test, a replicate pair was removed and tested on with a random forest classifier trained on the remaining 61 replicate pairs. This resulted in 62 “tests” with slightly different random forest classifiers. The importance of each feature is a metric of the influence that that feature has on the random forest outcome. The importance of each feature was extracted from each of the 62 tests and averaged. The average importance of each selected wavelength in the power spectrum was then plotted with the standard deviation. These data show that long wavelengths are highly important to the outcome (best differentiate power spectra of resistant and susceptible isolates), but the 0 frequency amplitude (long wavelengths) in the power spectrum correlates with the average height and therfore does not need to be added to our feature list. Instead we look at the next maximum importance at 18µm, which we include as a feature.

**Fig. 7.**
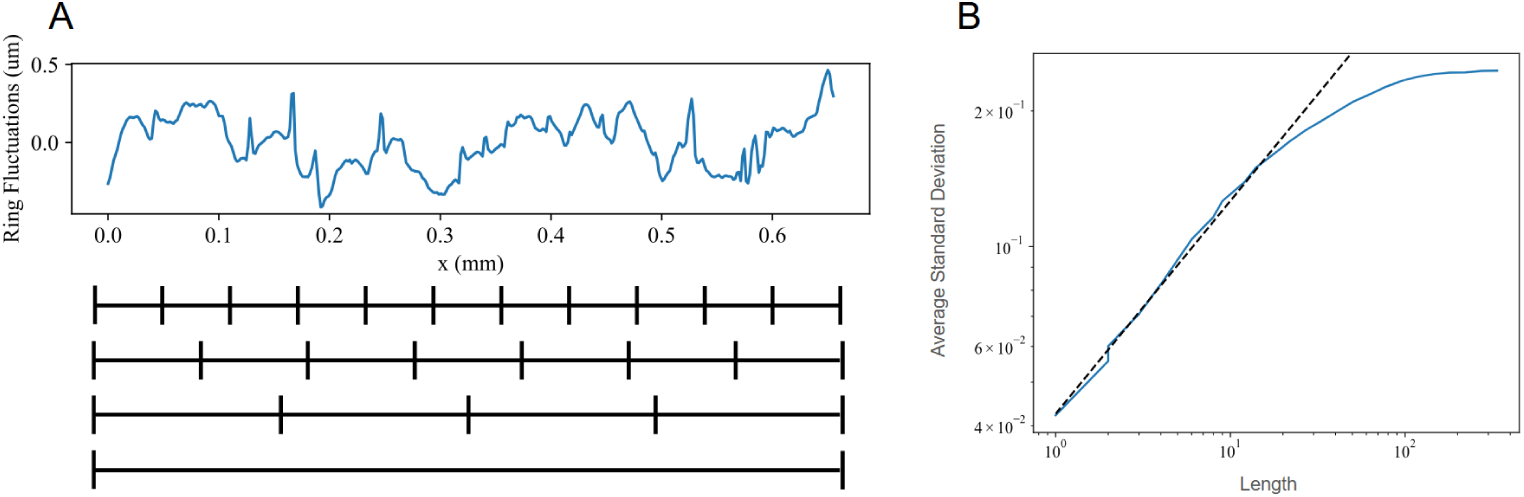
Defining the calculation of the Hurst exponent. **A**) The coffee ring profile is extracted by averaging the tallest pixels in rows perpendicular to the coffee ring. A line is fit to the ring profile and subtracted to leave only the fluctuations. The black dividers underneath represent varying bin lengths, *ℓ*. The standard deviation of each section is calculated and averaged across the fluctuation profile. **B**) These average standard deviations are called widths and plotted against the corresponding length, *ℓ*. The first linear region in the log-log plot is power-law scaling and the slope of that line, *H* (*w ∼ ℓ^H^*), is known as the Hurst exponent and is calculated as one of the features.

We also compare how initial cell density impacted dynamics by simply comparing the mean coffee ring and homeland heights. We consider the product of the heights, as well as the ratio between the two. The product is maximized when growth is observed in both regions, and minimized when both regions fail to grow, thus amplifying those individual signals when they match. The ratio of the height of the coffee ring to the height of the homeland characterizes how the initial distribution of cells impacts growth dynamics.

Finally, we include nine wavelengths from the power spectrum of the homeland, calculated by radially averaging the Fast Fourier Transform amplitudes. The nine wavelengths span the length scales of the image and are spaced logarithmically (in µm): 2.5, 5.2, 10.8, 22.5, 46.6, 96.6, 200.2, 415.1, and 860.5.

### 6.5 A two stage ML pipeline for calssifying experiments

As described in the main text, due to our limited statistics, we use LOOCV. Within the LOOCV, we used recursive feature elimination (RFE) as described in the methods to extract a subset of features most relevant to that classification. Samples were inoculated in duplicate to measure technical replicates. For LOOCV, we treat each replicate pair as a single entity to avoid potential data leakage. We remove both replicates from the training data, train on the remaining N-1 sample pairs, and then test both replicates. Thus the test strain is never used for recursive feature elimination or to train the classifier. We started with all features, and then performed RFE removing one feature each time and calculating the resultant accuracy. We will primarily focus on the number of features that maximized accuracy; when there was a tie, we went with the smallest number of features that maximized accuracy.

As we are data limited, we crafted a pipeline that proceeds across two stages. We first perform LOOCV and RFE on combinations of genera; one group is *Acinetobacter* and *Pseudomonas*, and the other is *Escherichia* and *Klebsiella*. Samples then proceed to a LOOCV and RFE between the two genera of their grouping. For example, if a sample is called *Escherichia* / *Klebsiella* in the first round, we would then perform another round of LOOCV and RFE just on *Escherichia* and *Klebsiella* to assign a predicted genera to each sample. We follow this protocol regardless of if the first round is correct or incorrect.

### 6.6 The simulation ML approach

A multi-head neural network classifier with a shared residual multi-layer perceptron trunk was trained via five-fold cross-validation to predict seven simulation parameters (*α*, *β*, L, D, fraction of cells initially in the coffee ring, number of clusters initially in the homeland, and the initial width of the coffee ring) from nine topographic features, using balanced class weights and softmax cross-entropy loss to handle uneven class distributions. The model architecture employed 16 residual blocks with layer normalization, GELU activations, and dropout, implemented in JAX/Flax and optimized with a cosine-annealed learning rate schedule.

### 6.7 Supplemental Figures

#### Supplementary information

**Fig. S1.**
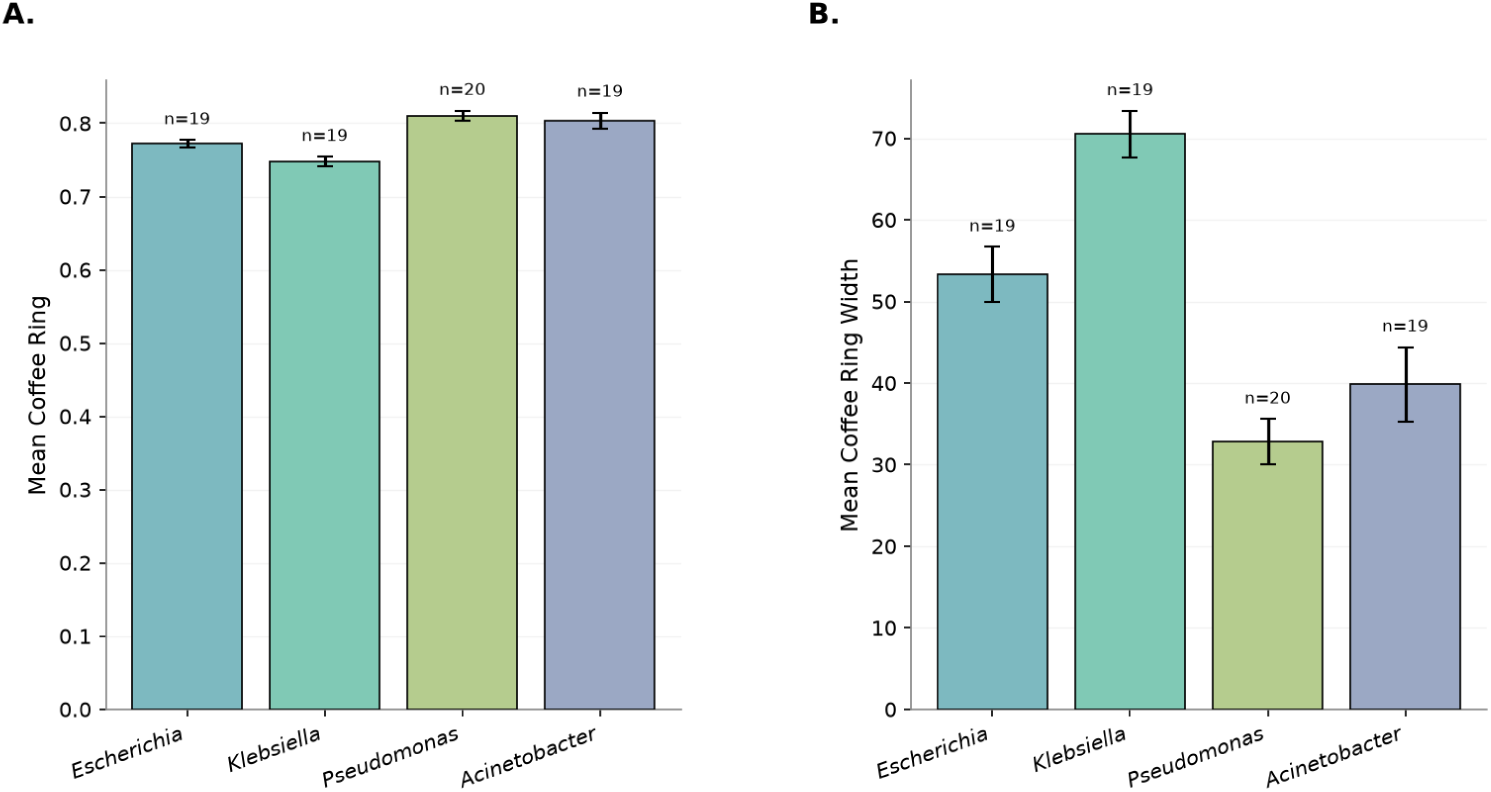
Coffee ring features vary by genera. A. The mean height of the coffee ring varies a small amount across the four genera we studied. B. The mean coffee ring width varies substantially. In both panels, error bars are standard deviations.

**Fig. S2.**
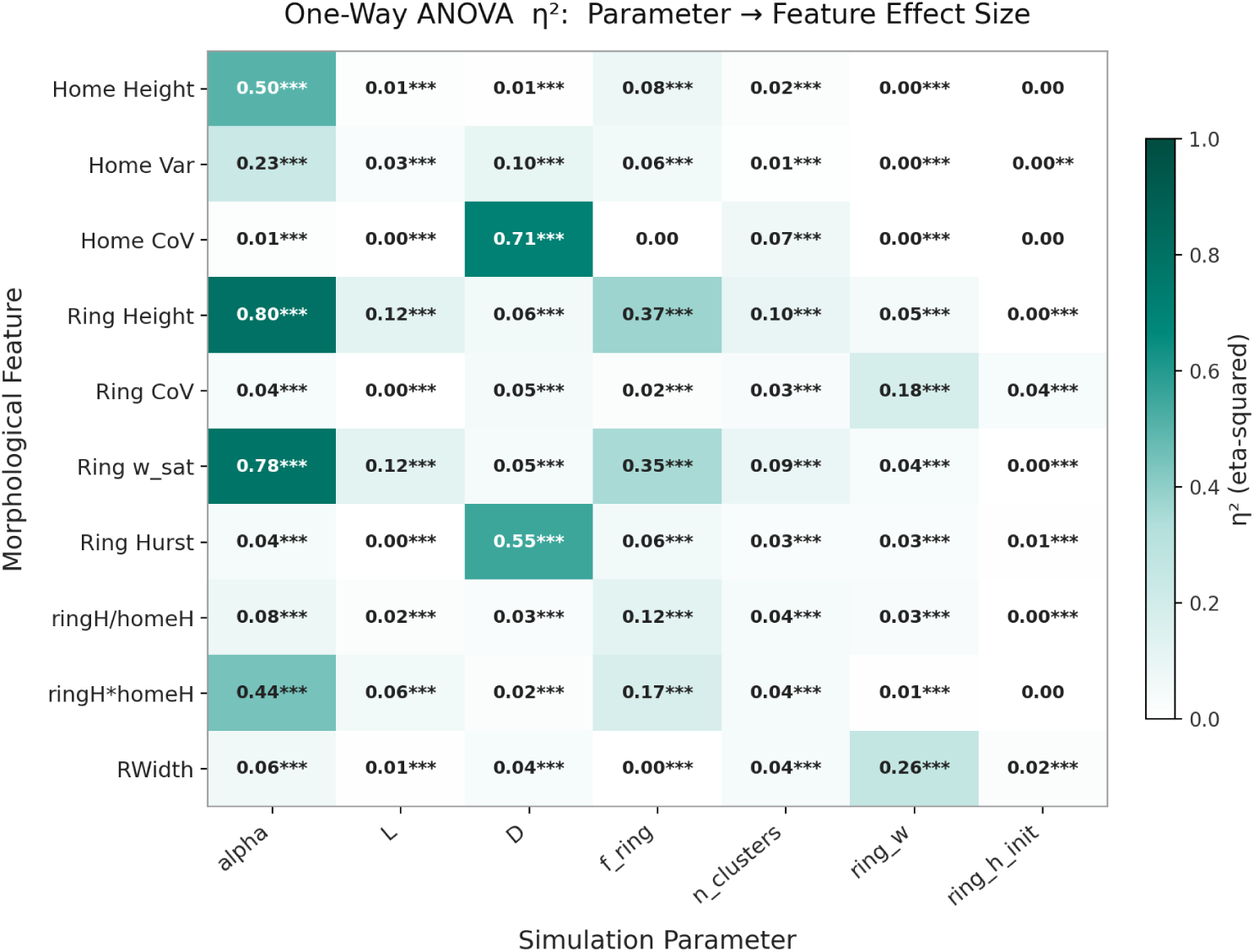
A heat map of *η*^2^ effect sizes from one-way ANOVA is shown, linking simulation parameters to simulation features.

**Fig. S3.**
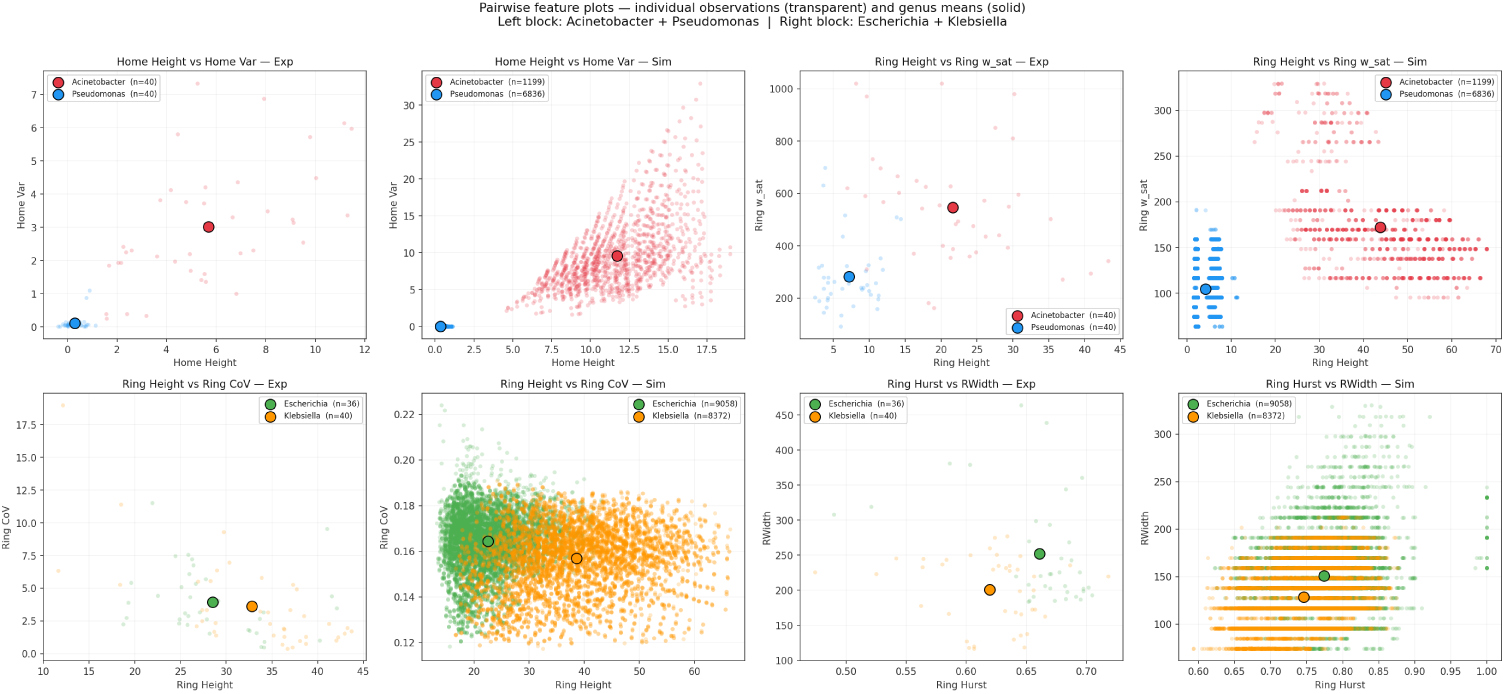
Simulation features and plotted against experimental features, for *Acinetobacter* and *Pseudomonas* (top row) and *Klebsiella* and *Escherichia* bottom row.

**Fig. S4.**
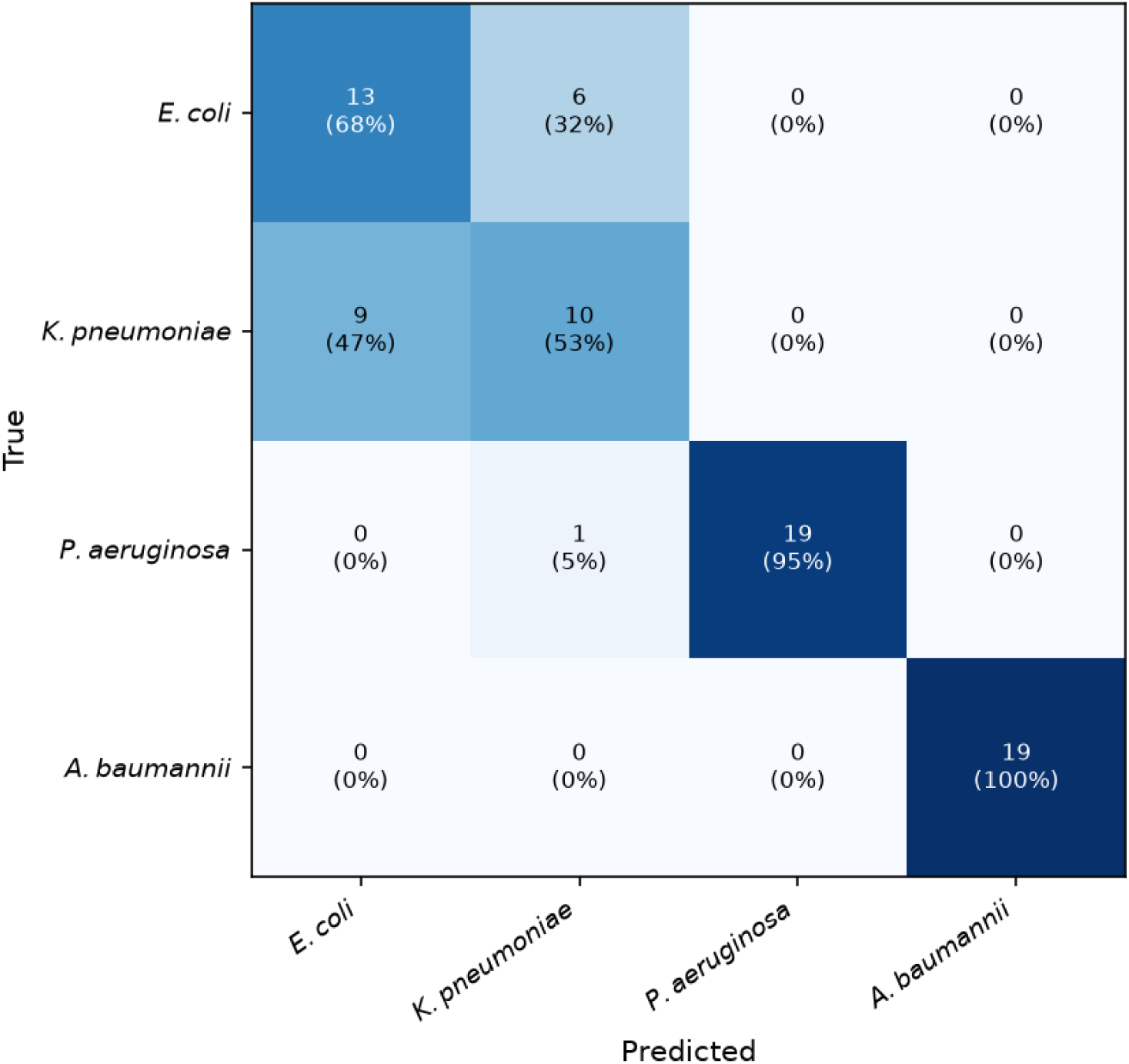
A confusion matrix when only a single parameter, the mean coffee ring height divided by the mean homeland height, is allowed for training.

**Fig. S5.**
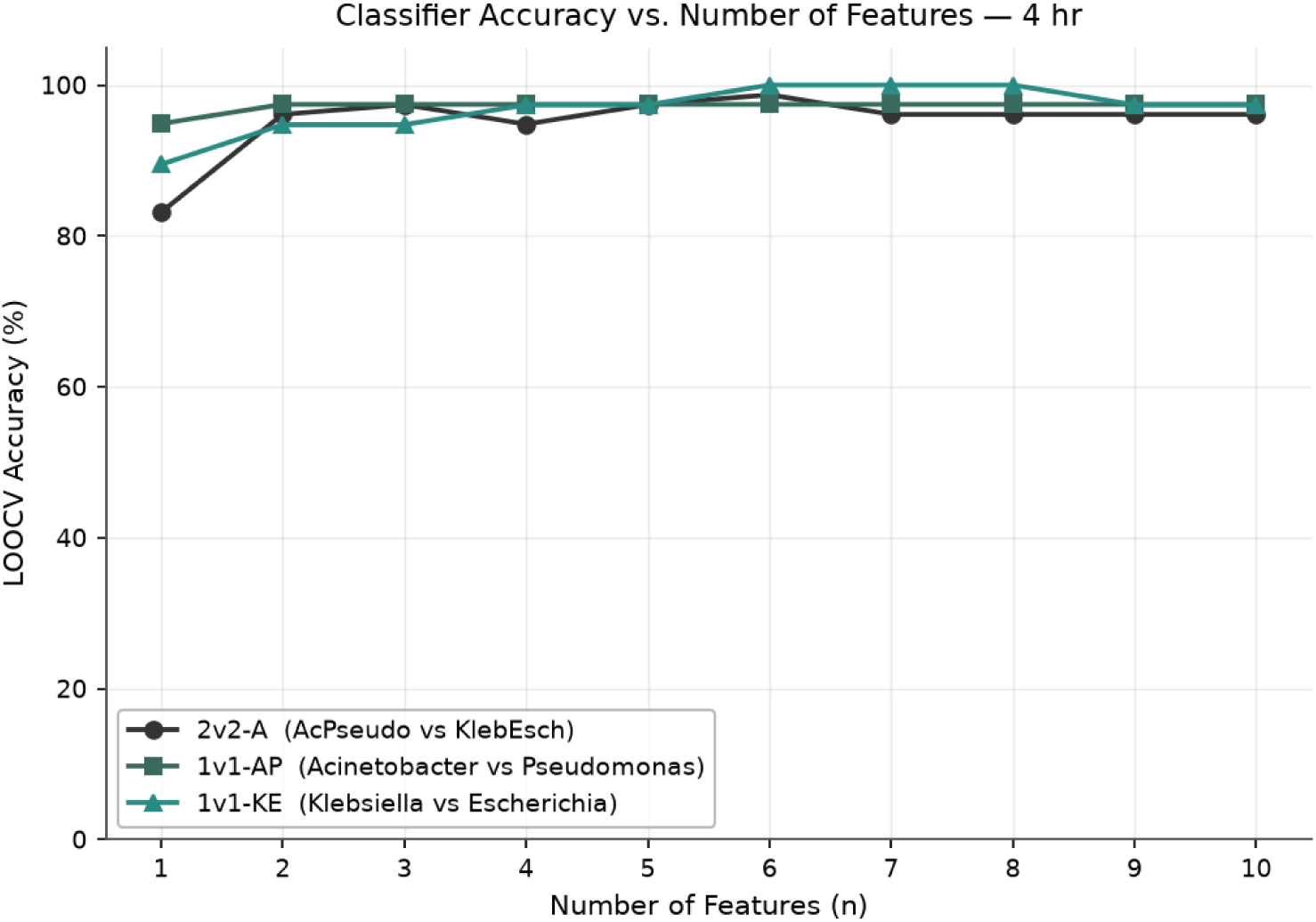
The accuracy of each of the three parts of the ML pipeline as a function of the number of features kept. While *∼* 6 features is typically optimal, accuracy is high for each part so long as the number of features is *≥* 2.

**Fig. S6.**
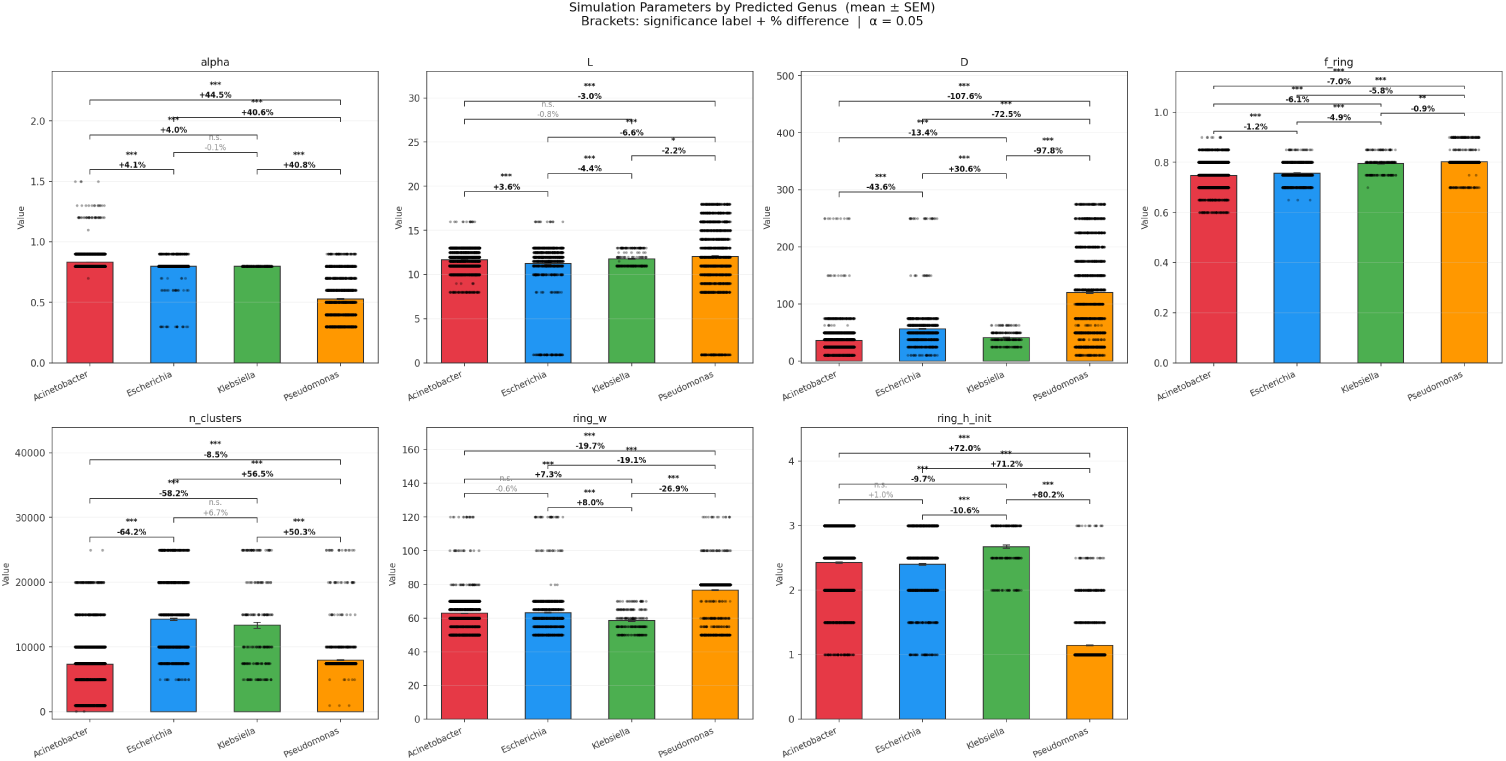
Summary of the properties of simulated topographies that correspond to different species. *∗ ∗ ∗* : *p <* 0.001; *∗∗* : *p <* 0.01; *∗* : *p <* 0.05; *n.s.* : *p ≥* 0.05.

#### Acknowledgements

We thank Sarah Satola for access to the strains used here.

## 7 Data and code

Data and code can be found at https://github.com/peter-yunker/Topography_ID_AST.

## Declarations

Drs. Krueger, Weiss, and Yunker are founders and equity holders in TopoDx, which is developing technology related to the research presented in this manuscript. TopoDx may benefit financially from the publication of these results.

Drs. Yunker and Krueger and Georgia Tech are entitled to royalties derived from TopoDx’s sale of products related to the research described in this paper. This study could affect his personal financial status. The terms of this arrangement have been reviewed and approved by Georgia Tech in accordance with its conflict of interest policies.

## Appendix A Section title of first appendix

